# Plasticity of genetic regulation during cellular learning

**DOI:** 10.64898/2026.05.27.728101

**Authors:** Lina Heistinger, Iuliia Parfenova, Aliaksandr Damenikan, Agnès Michel, Benoît Kornmann, Anna Sintsova, Yves Barral

**Affiliations:** Institute of Biochemistry, Department of Biology, ETH Zurich, Zurich, Switzerland; Department of Biochemistry, University of Oxford, Oxford, UK; Institute of Microbiology, Department of Biology, ETH Zurich, Zurich, Switzerland

## Abstract

Even under physiological conditions, cells are exposed to multiple, spatially and temporally complex signals and need to regularly adjust their response to survive and function. At least in some cases, identifying the appropriate response relies on the ability cells to learn from experience. For example, budding yeast cells learn to ignore futile mating signals. Using this paradigm, we investigated the genetic requirements supporting learning in single cells. We show that it arises from the dynamics of a large, highly redundant and plastic genetic network, where the contribution of most genes to learning varies across experimental replicates. Furthermore, this network is highly resistant to genetic perturbations and involves a broad panel of cellular functions and sensing pathways, suggesting that it can integrate a large diversity of inputs. Together, our data support the notion that cellular learning is an emerging feature of the information storage capability inherent to large sets of interacting genes, indicating that it should be widely conserved across cell types.

## Introduction

Cells are generally seen as responding to signals in a reproducible, pre-scripted manner. This conception is supported by many studies, establishing that cells respond to stimuli through specific, hierarchical pathways acting each downstream of a receptor or a sensor and eliciting highly reproducible effects. Such pathways have been characterized for a broad variety of stimuli, including stresses, nutrients and signalling molecules, such as hormones and morphogens^1–8^. These studies typically focus on well-defined stimuli at high and steady levels that promote unambiguous responses. However, under physiological conditions cells are exposed to diverse combinations of possibly contradictory inputs that are generally fluctuating in time and non-homogenous in space, with typically lower intensities than experimental ones. Thus, under physiological conditions, predetermined responses might not be appropriate. Rather, cells might need to make adaptive decisions that integrate different dynamic signals and generate outputs that protect their survival and if possible, their function. This would mean that cellular responses are plastic and “purposeful” rather than pre-scripted. However, how cells would achieve such purposefulness is largely unclear.

Interestingly, a variety of unicellular organisms and individual cell types in multicellular organisms show learning capacities that might shape purposeful responses. For example, unicellular eukaryotes as diverse as slime moulds, ciliates, and yeast demonstrate the ability to anticipate inputs in response to another that was temporally correlated in the past^9,10^, to become sensitized to consequential stimuli that they have recently encountered^11^, and to habituate to and hence, ignore perturbations when they are inconsequential^12–16^. These basal learning processes help prioritize decisions as a function of whether recurring inputs were informative, profitable, harmful or inconsequential in the past. Accordingly, theorical studies indicate that in varying environments the ability to store and use information about past events can provide strong fitness advantages to cells^17,18^. However, whether these integrative responses reflect truly plastic capabilities of cells to adapt their behaviour to diverse environments or are rather based on pre-scripted, hard-wired, higher-order control pathways is unknown. Any adequate understanding of cellular behaviours will first require a better knowledge of the mechanisms underlying their apparent plasticity.

To study this question, we use the budding yeast *Saccharomyces cerevisiae*, and its habituation-like ability to ignore futile mating signals^19^. Haploid yeast cells exist in one of two mating types, *MAT*a and *MAT*α. They signal their presence to cells of opposite mating type through pheromone (hormone-like peptides, called a- and α-factor, respectively). Upon binding to their cognate G-protein-coupled receptor (GPCR) on the cells of opposite mating type, these pheromones stimulate a mitogen-activated protein kinase (MAP-Kinase) signalling cascade^5,6^. At physiological pheromone concentrations (0.3-5 nM)^20^, the cells respond transiently to this signal, activate the expression of mating genes, arrest the cell cycle in its G1-phase and grow a mating projection, called a shmoo, towards the signalling partner. Reciprocally committing cells shmoo towards each other and eventually fuse to form a diploid zygote. The cells that fail to find a partner in due time stop shmooing, and resume proliferation. As cells learn and memorize that this pheromone signal is vain, they start ignoring the ongoing or any subsequent pheromone signal of similar level and keep this pheromone refractory state for most of their remaining lifespan^19^. Also, cells pre-exposed to short pheromone stimuli habituate to prolonged exposures more rapidly than unchallenged cells. Furthermore, habituated cells exposed to increased pheromone levels restart shmooing^19^. Thus, yeast cells modulate their response according to their memorized experience. Strikingly, this behaviour is not driven by receptor adaptation or desensitization; signalling from the receptor and the transcriptional effects downstream remain intact^19^. Given our extensive knowledge of the yeast pheromone response pathway, we reasoned that this learning-like response was a powerful paradigm for investigating the molecular mechanisms governing the apparent plasticity of cellular behaviours.

Thus, we sought to identify the genes that control habituation to futile pheromone exposure in *S. cerevisiae*. Particularly, we asked whether these genes function as a hierarchical and reproducible pathway overarching pheromone response or as a plastic information-processing network.

## Results

### A genetic screen for factors regulating habituation to mating pheromone

To identify the genes involved in pheromone habituation, we aimed to screen for mutations that specifically modulate the ability of cells to grow in presence of low, physiological pheromone levels. At saturating pheromone concentrations (25 nM pheromone and above), wild type cells barely habituate and cannot divide. Only mutations that abrogate pheromone signalling altogether enable cells to proliferate under these conditions. Accordingly, past screens carried out in this regime have identified 16 genes that, when mutated, abolish pheromone response altogether. These include the pheromone signalling genes, encoding the pheromone receptor and its signalling cascade, and four genes (*SIR* genes) controlling their expression^5,6^, which we generically refer to as the pheromone signalling genes. Under low pheromone conditions, however, wild type cells habituate and proliferate, albeit slower, such that mutants are difficult to spot. Therefore, in order to identify habituation genes, we needed to address two challenges. First, we needed to identify growth conditions at which all wild type cells respond to pheromone but habituate relatively rapidly, such as to prevent the out-competition of habituation mutants by those abrogating pheromone signalling altogether. Second, we needed a method to distinguish the mutants showing increased (accelerated habituation) or decreased proliferation (defective habituation) in presence of low pheromone levels, specifically, from the bulk of neutral mutations anywhere else in the genome that do not interfere with habituation.

To select optimal growth conditions, we measured the time individual cells take to resume budding when exposed to a range of pheromone concentrations. Haploid *MAT*a cells were placed on solid media containing defined concentrations of α-factor pheromone, ranging from 6 to 18 nM, and imaged by time-lapse microscopy (Figure 1A). At each of these concentrations, all cells initially arrested their cell cycle in G1 phase (as indicated by the nuclear Whi5-tdTomato signal) and started shmooing. Over time, cells aborted shmooing and started budding again, as expected^19^. In the next cell cycle, the experienced mother cells did not arrest in G1 phase again, indicating that they had habituated and remembered. In contrast, their daughter cells naively responded to pheromone, as described^19^. Importantly, the time cells needed to habituate increased with increasing pheromone concentration (Figure 1B). While all cells in the population resumed budding within 4 h at 6 nM pheromone, at 18 nM almost 70% of cells were still responding to the signal after 18 hours. Thus, yeast cells ignore weak signals faster than strong ones, consistent with habituation behaviours^13^. Based on these results, we selected 4 to 8 nM α-factor as a range of concentration for screening. To address the second challenge, we used SAturated Transposon Analysis in Yeast (SATAY) as a screening strategy^21^ (Figure 1C). Here, libraries of more than 300 000 cells each, with each cell carrying a unique, independent transposon insertion at a random position in their genome, were generated. These libraries sampled the genome at a resolution of one insertion every 46 nucleotide on average (about 50 independent alleles for a representative 2 kb gene). These libraries were amplified and then expanded further in the presence of 0, 4, 6 or 8 nM of pheromone, all other growth conditions remaining identical. To assess the reproducibility of the approach, three independent but virtually identical replicates were carried out in the same wild type strain. Importantly, these replicates were performed a few months apart. The positions of the transposons remaining after selective expansion were retrieved by PCR, sequenced and mapped on the genome (see M&M and below). Statistical analysis of insertion distribution and read numbers across the genome, comparing the selective to the non-selective condition (0 nM pheromone) indicated which genes are selected or counter-selected specifically in the presence of pheromone. For each gene, the deviation from the expected number of insertions and reads, as estimated from the 0 nM pheromone condition, was expressed in fold standard deviation (z-score). These z-scores report semi-quantitatively on the intensity of selection (negative z-scores) or counter-selection (positive z-scores) that pheromone applies on individual genes. The genes with decreased insertion and/or read numbers (negative z-scores) are required for growth in low pheromone and hence for habituation. We refer to them as habituation genes. The genes with increased insertion and read numbers promote pheromone arrest. Some of them are simply pheromone signalling genes. The others act in inhibiting or delaying habituation to pheromone. We refer to them as courtship genes (in reference to^22^). Genes that show insertion and read numbers close to expected (z-scores between-1 and 1) contribute neither positively nor negatively to habituation in our screening condition and are referred to as neutral genes.

**Figure 1:**
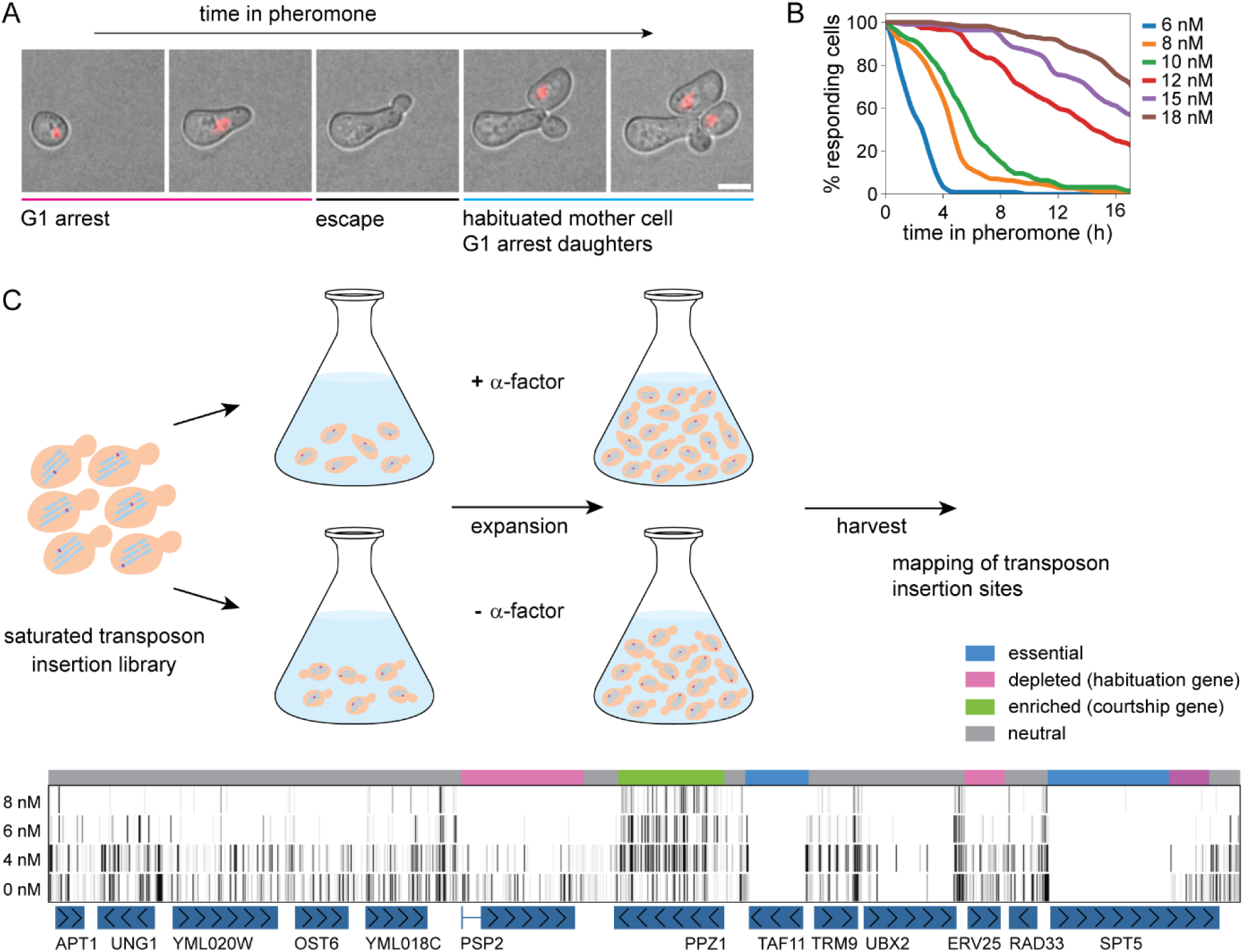
A genetic screen for factors regulating habituation to mating pheromone. **A)** Time-laps images of a haploid *S. cerevisiae MAT*a cells exposed to 12 nM α-factor on solid medium. The duration of cell cycle arrest (time to habituation) and the subsequent cell cycle profiles of mother and daughter cells are measured using cell morphology and the nuclear localization of Whi5-tdTomato as markers. Scale bar: 5 µM. **B)** Time to habituation in response to indicated α-factor concentrations as measured by time-laps microscopy. >50 individual cells per concentration. **C)** SATAY screening procedure. A library of cells carrying each a single transposon insertion at a random genomic position was split and expanded in the presence of 0, 4, 6 or 8 nM α-factor. After expansion, the insertion sites of the transposons remaining in each culture were mapped by sequencing. A representative region with mapped insertions (vertical lines) is shown. Comparison of the insertion pattern in the presence or absence of pheromone identify habituation (depleted in insertions), courtship (enriched in insertions), essential (no insertions) and neutral genes.

Analysis of our three replicates of the habituation screen clearly demonstrated that the concentration of pheromone had strong effects on the outcome (Figure 1C). The libraries selected at the lowest pheromone concentration (4 nM) had similar numbers of transposon insertions and sequencing reads per gene as the unselected libraries (Supplementary Figures 1A+B). In contrast, at 6 and 8 nM pheromone, increased selection drastically reduced the number of independent insertions, despite the total number of sequencing reads per library increasing. Furthermore, these reads mapped to fewer genes, strongly reducing the median number of reads per genes and thereby the dynamic range. At 8 nM pheromone, more than 40 % of all reads mapped to only ten pheromone signalling genes. At these concentrations pheromone insensitive mutants were taking over in the population, overshadowing habituation mutants. Therefore, we focused our analysis on the screens carried out at 4 nM pheromone.

### Habituation involves a large network of genes

Since our screen was replicated three times, we first analysed the results of each replicate separately, ranking the genes according to their z-score (see M&M, Supplementary Figures 2A-C). To compare the results across screens, we plotted the genes as a function of their z-scores in one individual replicates against another (Figure 2A, Supplementary Figures 2D-E). As expected, most genes fell along a broad diagonal, with the vast majority of them centred around 0 (−1.5 to +1.5). Also, a substantial number of the genes near the diagonal had lower or higher z-scores (≤-1.5 and >+1.5) in both plotted replicates, indicating that they showed reproducible changes in these two replicates. Indeed, a Wald test with Benjamini–Hochberg correction (see M&M), showed that 82 genes deviate significantly from neutral across all three datasets, indicating that they are consistent hits (Supplementary Figures 2F-G, adjusted p-value ≤0.05, orange and red dots in Figure 2A and Supplementary Figures 2D-E). Validating the selection procedure, the known pheromone signalling genes (red dots) were among the most enriched in insertions and reads, and the most significant across all datasets. Strikingly, however, many additional genes showed strong phenotypes in individual replicates (replicate-specific hits with at least one stringent z-score <-3 or ≥+3 are indicated as blue dots in Figure 2A and Supplementary Figure 2D-E). While most of these genes showed a strong z-score in one or two replicates, they showed neutral or even an opposite z-score in at least one of the other replicates. Therefore, they were not identified as statistically consistent when combining the replicates.

**Figure 2:**
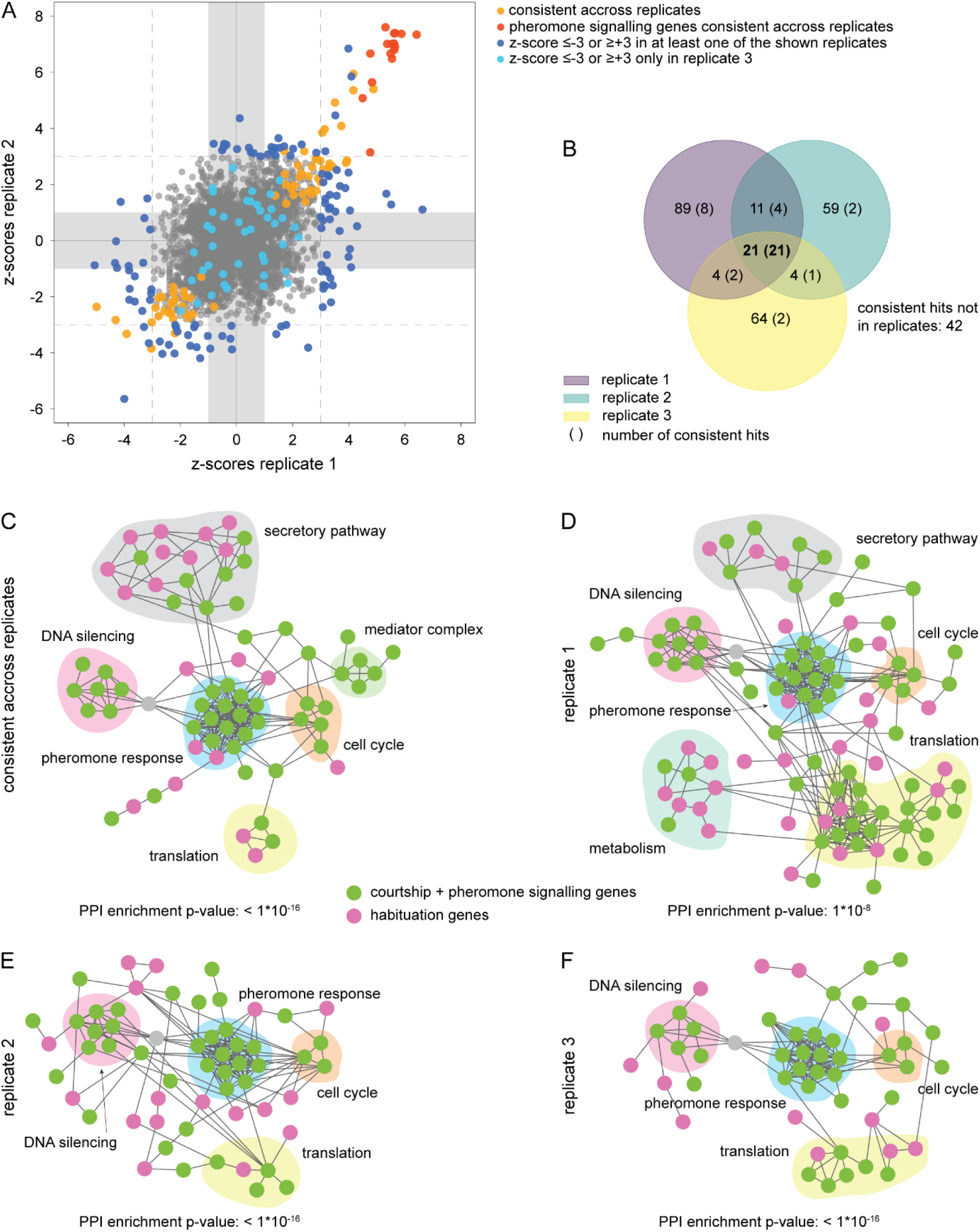
Hits identified in the individual screen replicates form dense networks of protein-protein interactions. **A)** All genes with a measurable z-score in replicates 1 and 2 are plotted as a function of their z-scores. Grey area: z-score values within one standard deviation. Dark blue dots: genes that pass the z-score threshold of ≤-3 or ≥3 in at least one of the two replicates. Light blue dots: genes that pass the threshold of ≤-3 or ≥3 only in the third replicate. Orange dots: statistically significant (consistent) genes across all three replicates. Red dots: Expected silencing and pheromone signalling genes consistent across all three replicates. Grey dots: Neutral in each replicate and across the three replicates. **B)** Venn diagram showing the overlap between gene hits identified in the individual replicates (replicate 1: 125 genes, replicate 2: 95 genes, replicate 3: 93 genes; z-score ≤-3 or ≥+3). The number of consistent hits across replicates within each group (82 genes from combined statistical analysis; adjusted p-value ≤0.05) is shown in brackets. All 21 hits identified in all three replicates are consistent hits. **C)** Predicted protein-protein interaction network for the hits consistent across replicates (82 genes, adjusted p-value ≤0.05). The predicted protein-protein interactions between products of the gene hits were calculated and compared to the number of interactions observed in similar size sets of random genes using STRING. *HML*α (grey) is added as anchor. The main fully connected network (70 nodes) is shown. Courtship and pheromone signalling genes are plotted in green, habituation genes in magenta. **D)-E)** Predicted protein-protein interaction network for the hits with z-score ≤-3 or ≥3 in the individual replicates (replicate 1: 125 genes, replicate 2: 95 genes, replicate 3: 93 genes). The predicted protein-protein interactions were calculated as in C). *HML*α (grey) is again used as an anchor. As in C), only the main fully connected networks are shown (replicate 1: 100 genes, replicate 2: 58, replicate 3: 50). Colours as in C).

A striking feature of these replicate-specific hits is that in at least one replicate, their z-score deviated much further from zero than those of the consistent hits (see blue vs orange dots in Figure 2A), suggesting that they may have strong phenotypes. Alternatively, they could correspond to experimental noise. Intrigued by the first possibility, we sought to test whether they were false positives or true hits. One indication that the hits were not random came from the SATAY screen itself. All hits are identified by many independent alleles corresponding to distinct individual transposon insertions, and thus, by several biological replicates in the pool. These distinct alleles thereby confirmed each other within each replicate.

Next, we reasoned that if hits were false positives, they should correspond to random genes, whereas if they were true hits, they may share some non-random common features or function in common cellular processes. To investigate this question, we asked whether the products of these replicate-specific hits were more likely to physically interact with each other and with the products of the consistent hits than expected by random. Therefore, we performed network analysis based on known protein-protein interactions using STRING^23^. For each replicate, this analysis was restricted to all the genes with the most extreme z-scores (stringent z-score cutoff of ≤-3 or ≥+3; ranging from 93 to 125 genes in the different replicates). In parallel, the same analysis was performed with the 82 hits identified as statistically consistent across the three replicates, of which 42 do not pass our z-score cut-off in any replicate. The overlap between these four analyses comprised only twenty-one hits (Figure 2B), which included all the pheromone signalling genes. Remarkably, these analyses showed a strong and highly significant enrichment of interactions within each dataset, connecting many of the identified hits, including replicate-specific ones, into extended interaction networks (Figure 2C-F; p<10^-8^). As expected, these networks reproducibly included functional clusters involved in pheromone signalling (*STE* genes, *GPA1*, *FAR1*, *DIG1*), cell cycle (*WHI3*, *WHI4*, *WHI5*, *CLN3*) and DNA silencing (*SIR* genes, *NAT1*). Some functional clusters, including the mediator complex and the secretory pathway, were strongly visible in the network of consistent hits (Figure 2C) but were smaller or absent from the networks identified at stringent z-score cutoff in the individual replicates. This suggests that these functional modules have consistent but moderate effects on habituation. Strikingly, the opposite was observed for a cluster involved in translation, which was strongest in the network based on individual replicates. Interestingly, a cluster of ten genes associated with carbon and amino acid metabolism was above our z-score cutoff only in one screen (Figure 2D). Overall, however, all datasets produced very similar protein-protein interactions networks, highlighting that they globally comprised similar functionalities. The networks based on the individual replicates linked a large fraction of the consistent and replicate-specific hits to each other. Thus, many if not most replicate-specific hits were not random, but part of the same protein-protein interaction network. This suggested that many genes engaged in habituation sporadically across replicates. Therefore, we will refer to these hits as sporadic.

To test this hypothesis more rigorously, we next combined all 252 hits above cutoff in at least one replicate and the 82 consistent hits (294 genes total) and asked whether they were indeed part of the same network. Remarkably, this analysis predicted a highly significant network protein-protein interaction pan-network (Figure 3) of 251 connected nodes (85% of the input genes, p<10^-16^). The 21 genes identified in all datasets (7 % of all hits, red labels) were all enriched in transposon insertions, consistent with their role in pheromone signalling. Additional 61 genes (20.5 % of total) were consistent hits, 19 of which (6.5 %, orange labels) were also above cutoff in at least one replicate and 42 were moderate hits across replicates (yellow labels). Thirteen genes (4 %; light blue) were sporadic hits above cutoff in two out of three replicates. However, the vast majority of genes (200 genes, 68 % of all hits and 80% of all nodes) were sporadic hits identified in only one replicate (dark blue), that connected into distinct functional clusters. Thus, we concluded that at the chosen cutoff only very few if any of the sporadic hits were false positive. We concluded that habituation relies on the activity of an extended gene network. Furthermore, the weight of many of the nodes in that network varies between replicates, indicating that the habituation network is plastic.

**Figure 3:**
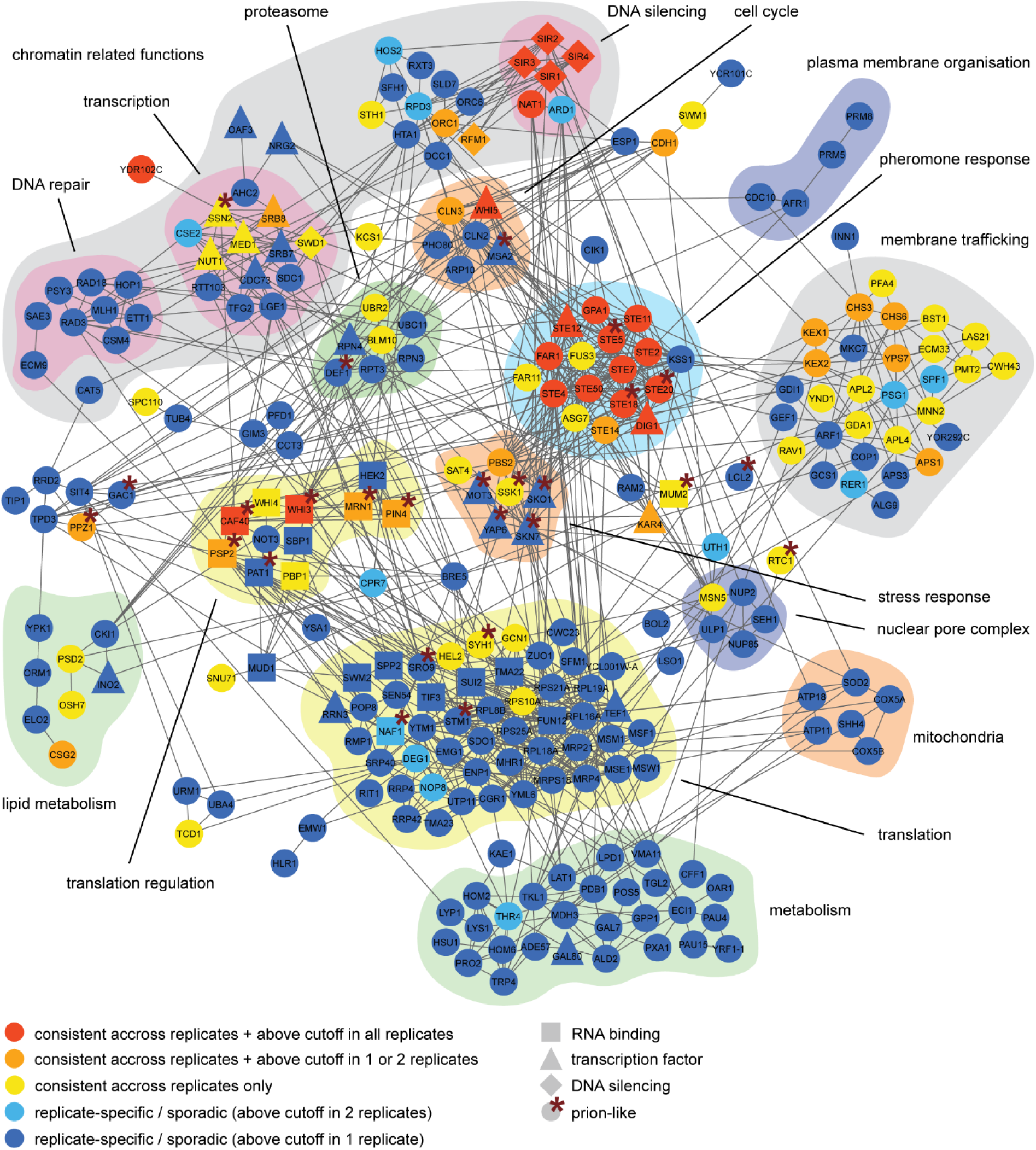
The pan-network of all habituation hits connects many different cellular functions. Predicted protein-protein interaction network of all hits identified in the individual screens (at cutoff of z≤-3 or ≥+3) and the combined statistical analysis (adjusted p-value ≤0.05). 294 hits in total resulted in a network of 251 connected nodes using STRING (enrichment p-value <10^-16^). Five additional genes which were just below read / transposon number abundance cutoff (*PRM5*, *SBP1*, *NOT3, CSE2*) or only identified as a hit in the *whi5*Δ strain (*KSS1*) and affected habituation in our validation experiments (see Figure 5B) were also included. Hits were manually ascribed to functional clusters. Node colour and shape code is indicated within the figure.

Inspection of this pan-network highlighted the diversity of cellular functions and structures involved in pheromone habituation. We anticipate that some of the genes, such as those involved in stress signalling, DNA repair and metabolic function (lipid and amino acid synthesis, carbon catabolism and mitochondrial function), convey input signals to the network. The DNA silencing cluster is known to control mating-type identity, another form of input. On the other hand, other genes probably deliver network outputs, enacting habituation. This is the likely role of the cell cycle regulators. Remarkably indeed, all cell cycle genes identified in our screen act at the G1/S transition. Other functional clusters present in the network are less easily classified as input or output and might therefore function in the computing events that integrate and store information and make decisions. The chromatin-associated cluster (41 genes, 14% of all hits, including 22 transcription factors, 6 components of the Mediator complex and 13 histone modification factors) are likely to contribute to computing. Genes involved in translation and translation regulation are also good candidates for processing information. This cluster includes RNA-binding proteins involved in mRNA stability and translation regulation (11, 4 %), tRNA modification, ribosome assembly and ribosomal subunits (translation cluster of 50 hits, 17 %). However, some functional clusters may act simultaneously in input collection, information processing and output signalling. The cluster of 30 genes involved in membrane trafficking may be one of them. Particularly, the Golgi proteases Kex1 and Kex2 have known roles in exocytosis and processing of secreted proteins, including the α-factor pheromone. Remarkably, the cells in which the screen was carried out do not secrete α-factor but habituate to it. Thus, Kex1 and Kex2 may function both in pheromone production in one partner and modulating the response to pheromone, i.e., habituation, in the other partner, if not in both. Thereby, these molecules are excellent candidates for linking signalling and response. Thus, our data indicate that the pheromone habituation network comprises and connects many cellular functions and possibly integrates many signals beyond pheromone alone. Furthermore, the high level of connectivity suggests that the network may comprise many feedback loops and may therefore function in a recurrence-based manner.

In this respect, the network contains four classes of proteins of particular interest: 1) the regulators of chromatin organization (see above), 2) transcription factors (see above), 3) RNA-binding proteins involved in mRNA stability and the regulation of their translation (19 / 6% of hits) and 4) proteins with prion-like domains or behaviours (26 / 9%, marked with * in Figure 3). The function of these genes in the regulation of gene expression makes them excellent candidates for regulating each other and for functioning in cellular memory.

Interestingly, sporadic and consistent hits showed distinct distributions in the network. All strong and consistent hits fall into only few clusters, particularly those involved in pheromone signalling (Figure 3, red nodes), except for *YDR102C* where insertions into its coding sequence may affect expression of the neighbouring pheromone response gene *STE5*. Genes with a moderate but consistent effect (orange and yellow) were particularly prominent in membrane trafficking and transcription. In contrast, the metabolism, translation and DNA repair clusters comprised mostly sporadic hits (blue). These different distributions support the notion that the different clusters contribute distinctly to network function.

Together, the data demonstrated that habituation to pheromone involves more than 250 genes, more than 10-fold the number of genes initially required for pheromone signalling. These genes function in a broad range of cellular processes and structures. They were not specifically associated with the pheromone response before and may function in integrating a broad variety of signals for the cell to decide whether to habituate to pheromone or not, probably in a context-dependent manner.

### Epistasis analysis confirms network behaviour and identifies additional habituation genes

Thus, our observations suggested that habituation to pheromone is controlled by a distributed, interconnected regulatory network rather than a linear pathway. To test this hypothesis, we next performed epistasis analyses with five genes in the network, asking whether their function depends on each other, as expected in a pathway, or whether they function independently of each other in habituation, as expected from a network structure. These five query genes, *WHI3*, *WHI5*, *PIN4*, *PPZ1*, and *CAF40*, were all consistently enriched in transposon insertions in the initial screens. Whi3 and Whi5 function in the control of the G1/S transition of the cell cycle. Whi3, Pin4 and Caf40 are RNA-binding proteins, and their products contain both N- and/or Q-rich domains typical of prion-like domains. Whi3 has been implicated in cellular memory already^19^. *PPZ1* encodes a phosphatase located at the plasma membrane and regulates potassium transport. For our analysis, each of the selected genes was deleted individually to generate five *MAT*a query strains. To extend the analysis beyond the interplay between these five query genes to their relationship to the rest of the network, we tested epistasis using the SATAY approach again. Indeed, we reasoned that if a gene X still scores as a hit in a genetic background where a query gene Y is deleted, then these two genes act independently of each other. Thus, we carried out new SATAY screens in our five query strains as described above.

Consistent with their identification as genes modulating pheromone habituation, all query strains responded effectively to pheromone but habituated faster than wild type, as determined by time-lapse microscopy (Figure 4A, Supplementary Figures 3A-E). For each mutant, a transposon mutagenesis library was generated, which was again grown under pheromone selection. At 8 nM pheromone the mutants habituated and grew comparably to the wild type at 4 nM, consistent with their fast habituation phenotype. Accordingly, sequencing indicated that for all five mutant strains the selection at 8 nM was optimal for analysis. Therefore, these datasets were processed and analysed as described above (Supplementary Figures 3F-J), using a single replicate for each strain.

**Figure 4:**
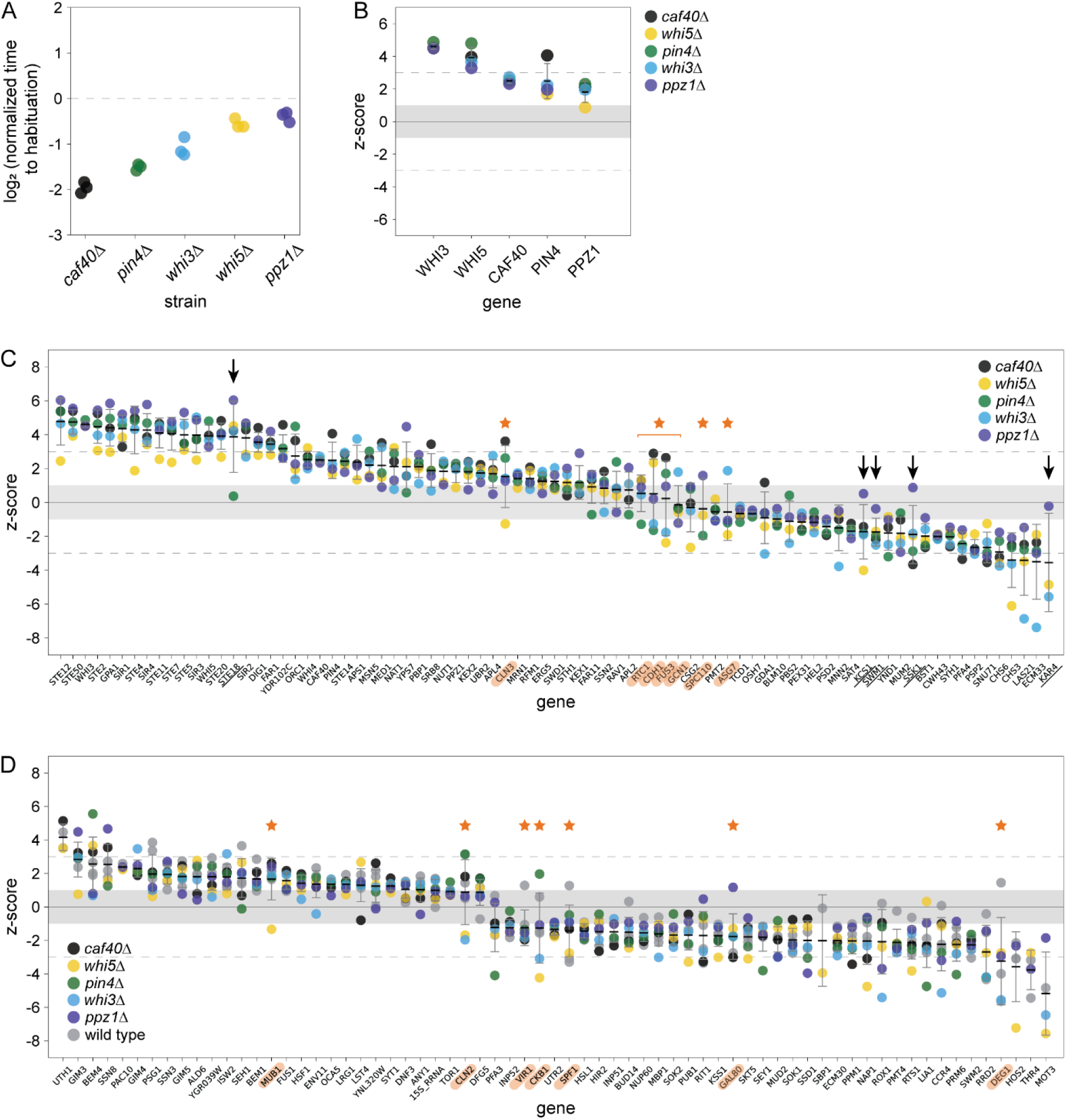
Screens with habituation mutants confirm network structure. **A)** Normalized time to habituation of individual query gene deletion strains. The duration of shmooing was determined by time-laps microscopy in the presence of 12 nM α-factor. For each mutant, three biological clones were analysed and normalized to the median habituation time of the wild type control imaged in the same experiment. Normalized median times calculated from 50 individual cells per sample are shown. For all mutants, time to habituation was significantly different from the wild type as determined by a two-sided Mann–Whitney U test with Benjamini-Hochberg correction (p-values <0.05). **B)** Z-scores of query genes in epistasis screens carried out in strains lacking one of the other query genes. Grey area: z-scores within one standard deviation. Individual values, median and standard deviation across screens are shown. The genotype of the strain in each epistasis screen is indicated by the dot colour. **C)** Z-scores of the 82 consistent hits (across wild type replicates) in the five epistasis screens. Genes are ranked by their median z-scores. Median, error bars, and grey area as in B). Arrows: genes that become neutral in one of the mutants. Orange stars: bivalent genes. **D)** Z-scores of the 66 additional significant hits identified as consistent across all wild type and mutant replicates. The *ROX1* gene is added for illustration. Gene ranking, grey area, orange stars and error bars as in B) and C).

Remarkably, these epistasis screens established that all five query genes acted independently of each other in habituation: their z-scores were high in all other query strains (Figure 4B). The only exception was *PPZ1*, which scored just below 1 in the *whi5*Δ deletion strain (z-score = 0.9). In this strain, most genes scored lower than the average z-score over all mutant strain (Figure 4C), indicating reduced selection. However, the *WHI5* gene scored above cutoff (z-score = 3.3) in the *ppz1*Δ mutant strain. Thus, among all habituation hits, at least the Whi3, Whi5, Pin4, Ppz1 and Caf40 proteins contribute to habituation in the absence of each other, and hence, do not require each other for function. This indicated that they represent either five independent pathways or five independent nodes in a broader habituation network. In either case, these data demonstrate that the habituation network does not function as a linear pathway.

To gain deeper insights in the functional organization of this network, we filtered the results of the individual screens using the same strict z-score cutoff of ≤-3 or ≥+3 as for the wild type. We then analysed the density of protein-protein interactions within these lists of hits. All individual datasets were of similar size and strongly enriched in protein-protein interactions, comparable to the wild type datasets (Supplementary Figures 4A-E). The predicted network obtained with the *caf40*Δ strain was the smallest, likely due to a reduced number of sequencing reads resulting in the lowest number of hits compared to the other datasets. All five protein-protein interaction networks were structurally similar to each other and to those identified in wild type, highlighting the same key cellular processes, such as pheromone response, cell cycle progression, transcriptional regulation, osmotic stress response and translation. At the individual gene level, we observed the same sporadicity across the epistasis screens as between the wild type replicates. At our z-score cutoff, again only the pheromone signalling genes were hits across the five mutant screens. Thus, the network underlying pheromone habituation maintained the same general organization despite inactivation of any of our query genes. Thus, none of these five mutants abrogated any of the main components of the habituation network. We conclude that this large, distributed network functions in a highly redundant and robust manner.

This notion was further supported by analysing how the role of the 82 genes identified as consistent hits in wild type was affected in the query strains (Figure 4C). Remarkably, for virtually all these genes none of the query mutations affected their z-score trend. Only very few occasional genes became neutral in one of the five mutant background (black arrows, Figure 4C). For example, the pheromone signalling gene *STE18* was no longer enriched in insertions and reads in the *pin4*Δ mutant context. Careful analysis of the insertions mapping in this gene showed that the second half of the gene and its 3’UTR were depleted of insertion in that context, specifically. Thus, this sequence may be required for pheromone habituation specifically in the absence of the RNA-binding protein Pin4, potentially at the transcript level. Also, the Swm1 E3 ubiquitin ligase, a component of the anaphase promoting complex/cyclosome (APC/c), the phospho-relay and osmo-sensor protein Ssk1, the inositol hexakisphophate kinase Kcs1, and the transcription factor Kar4 were no longer promoting habituation in cells lacking the protein phosphatase Ppz1. Interestingly, for *SWM1* and *KAR4* this was caused by a strong reduction of insertions already at 0 nM pheromone in this mutant, suggesting that these genes are synthetic lethal with *PPZ1*, indicating that they function in parallel pathways. In contrast, Ssk1 and Kcs1 may require Ppz1 function for habituation, putting them in a Ppz1-dependent subnetwork.

Interestingly, 7 of the consistent genes (orange stars, Figure 4C) scored as courtship genes in some query strains and as habituation genes in others, indicating that they acted in a bivalent manner in habituation, depending on the genetic context. For example, the MAP kinase Fus3, which propagates pheromone signalling in response to receptor stimulation, scored as a courtship gene in the wild type, *caf40*Δ, *pin4*Δ and *ppz1*Δ mutant strains and as a habituation gene in the *whi3*Δ and *whi5*Δ mutant strains. This indicated that Fus3 activity promotes both pheromone response and habituation and while the first activity limits cell growth in most genetic backgrounds, the second activity promotes it in cells lacking Whi3 or Whi5 functions. Similarly, *CLN3* was a courtship gene in most genetic backgrounds but a habituation gene in the *whi5*Δ mutant cells. In the *CLN3* gene, the region coding for the so-called PEST domain, which targets the protein for degradation by the ubiquitin-proteasome pathway, was enriched in insertion upon selection in pheromone, likely-resulting in higher Cln3 levels, whereas the so-called cyclin box, through which Cln3 activates the kinase Cdk1 and promotes cell cycle entry, was depleted of insertions (Supplementary Figure 4F). Whereas the insertions in the PEST domain dominated in most genetic contexts, the depletion of the cyclin box dominated in the *whi5*Δ mutant. Genes showing similar bivalent effects include the genes *RTC1*, coding for a positive regulator of the TORC1 kinase complex, *CDH1*, encoding another subunit of APC/c, *GCN1*, coding for an activator of the Gcn2 kinase (which was not a hit), and *ASG7*, coding for a regulator of Gβ in *MAT*a cells, specifically.

Together, these studies show that only few of the 82 reproducible hits require the function of any of the five query genes to promote or inhibit habituation. By extension, these data support the notion that the vast majority of them function independently of each other rather than as one or few linear genetic pathways. Thus, habituation to pheromone signalling appears to be the emergent response of a large, distributed, non-linear gene network. The bivalent genes identified in this network might be directly involved in switch-like decisions during habituation. The fact that they typically coded for key regulators of fundamental cellular events such as cell cycle entry (*CLN3*, *CDH1*), cell growth (*RTC1*, *GCN1*), and pheromone response (*FUS3*, *ASG7*) supports this conclusion.

We next reasoned that since the networks identified in the wild type and mutant strains are very similar, including all eight data set in our statistical analysis would increase our statistical power and potentially identify more of the genes acting consistently in habituation. Indeed, this approach identified 129 genes, of which 66 genes had not been identified in the previous statistical analysis of the wild type screen (Figure 4D). Among them, eight (*GAL80*, *GIM3*, *MOT3*, *RIT1*, *SWM2*, *CLN2*, *SEH1*, *RRD2*) were sporadic hits in the wild type screen. Many of the new hits encoded additional components of complexes already found in the wild type replicates, such as the prefoldin complex (*GIM4* and *GIM5*), the mediator complex (*SSN3* and *SSN8*), protein phosphatase 2A complex (*RTS1*, *RRD2*, *PPM1*), the P-body and associated factors (*SBP1*, *CCR4*, *PUB1*), RNA editing (*VIR1*), the TORC1 complex and its regulators (*TOR1*, *LST4*), and the osmotic stress response pathway (*MOT3*, *ROX1*, two transcription factors comprising prion domains). This approach also recovered the MAP-kinase gene *KSS1* as a mild habituation gene. Kss1 functions partially redundantly with the MAP kinase Fus3 in pheromone response, particularly at low pheromone concentrations, and is among the few factors previously known to function in recovery from pheromone-induced cell cycle arrest^24,25^.

Among the completely new cellular components emerging as consistent hits in this extended analysis, several genes encoded proteins involved in the formation of the mating projection and cell fusion upon mating (*BEM1*, *BEM4*, *FUS1*, *DNF3*, and *LRG1*). These genes all behaved as courtship genes, suggesting that the tip of the mating projection contributes to inhibiting habituation and that cell polarity may contribute to habituation decisions. Together, our epistasis analysis confirmes that habituation to pheromone is controlled by a large, highly distributed and plastic regulatory network.

### Network mutants show high phenotypic diversity but maintain ability to habituate

To gain more concrete insights into how the identified genes affect habituation and to further test our interpretation that even the sporadic hits contribute to this process, we characterized the habituation phenotype of 32 hit genes, comprising sporadic (14 genes) and consistent hits (18 genes) across habituation and courtship genes (11 and 21, respectively, Supplementary Figure 5A). We added four control genes (*GUT1*, *GAL3*, *CLN1*, *CAF4*), which remained neutral in all our screens. The entire open reading frames of these 37 genes were individually deleted, and the mutant cells were imaged as they responded and habituated to 12 nM pheromone (where habituation is slow, giving high sensitivity for analysis of habituation defects) over a period of 18 hours. From these data we recovered five parameters: Shmooing frequency in pheromone, time to habituation (defined as resumption of budding), memory defects (re-shmooing after habituation), inheritance of memory (daughters that fail to shmoo) and ability to modulate habituation time as a function of pheromone concentration. Images of cells presenting these different phenotypes are shown in Figure 5A.

**Figure 5:**
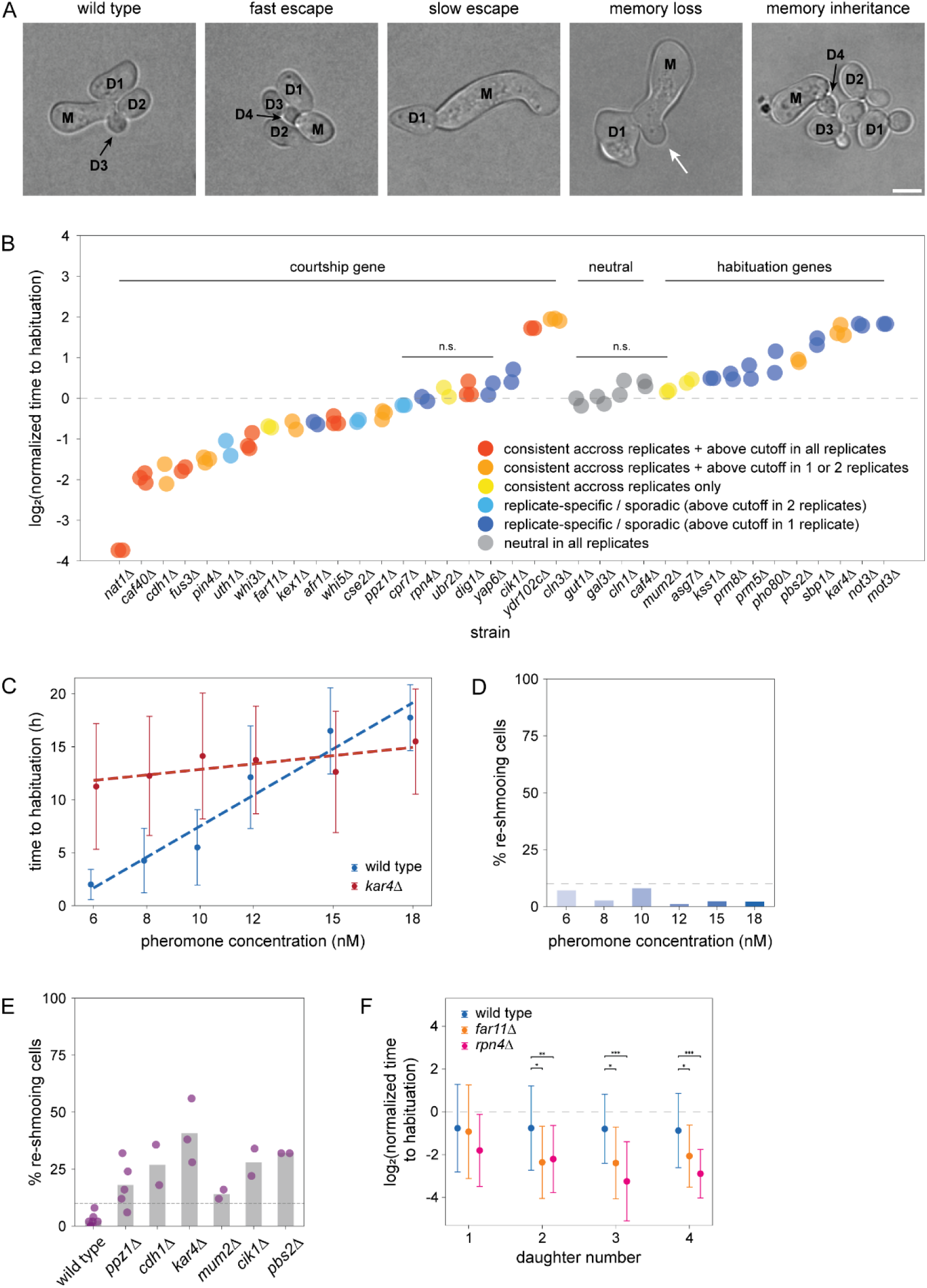
Network mutants show high phenotypic diversity. **A)** Representative bright field images of the observed habituation phenotypes, compared to wild type. M indicates the mother cell, D1-D4 indicates the daughters born by the habituated mother cell. White arrow highlights site of re-shmooing as indicator of memory loss. All cells were imaged on agar pads containing 12 nM alpha factor. Scale bar: 5 µM. **B)** Normalized time to habituation in individual network gene deletion mutants and control strains, measured and plotted as in Figure 4A. For each mutant, 50 cells of at least two independent clones were analysed. Consistent and replicate-specific hits and neutral control genes are shown in the indicated colours. For all mutants except the strains marked as n.s., time to habituation was significantly different from the wild type as determined by a two-sided Mann–Whitney U test with Benjamini-Hochberg correction (p-value <0.05). **C)** Median habituation time of wild type and *kar4Δ* mutant cells at indicated α-factor concentrations. The duration of shmooing was measured as in Figure 4A (wild type: 6 nM: 85 cells, 8 nM: 114 cells, 10 nM: 50 cells, 12 nM: 82 cells, 15 nM: 88 cells, 18 nM: 94 cells; kar4Δ: 6 nM: 98 cells, 8 nM: 80 cells, 10 nM: 98 cells, 12 nM: 150 cells, 15 nM 66 cells, 18 nM: 80 cells). Error bars represent standard deviation. Linear trend lines were added for improved visualization (R^2^ wild type: 0.95, R^2^ *kar4*Δ: 0.59). **D)** Fraction of re-shmooing wild type cells at indicated α-factor concentrations. Bars represent the percentage of habituated cells with at least one prolonged G1 arrest of 60 minutes or more. (6 nM: 85 cells, 8 nM: 114 cells, 10 nM: 50 cells, 12 nM: 82 cells, 15 nM: 88 cells, 18 nM: 94 cells) **E)** Fraction of re-shmooing cells in individual network gene deletion mutants at 12 nM α-factor. Bars represent the median fraction of re-shmooing cells measured as in D). Data points show the fraction of re-shmooing cells in individual replicates of 50 cells. All mutants with more than 10 % re-shmooing cell are shown. **F)** Habituation time in daughter cells born by habituated wild type, *far11Δ* and *rpn4Δ* mutant mother cells. Shmooing duration of the mother cells and their first four daughter cells was determined by time-laps microscopy in the presence of 12 nM α-factor. Habituation times of daughter cells were normalized to their respective mother cell. Data points represent median values of 27 to 35 cells per strain. Error bars represent standard deviation. Differences in normalized time to habituation between strains were analysed using a two-sided Mann–Whitney U test (* p-value <0.05, ** p-value <0.01, *** p-value <0.001).

All tested mutants responded to the pheromone signal, demonstrated by a prolonged residence time of Whi5 (tagged with tdTomato) in the nucleus and shmoo formation, indicating that none of them affected pheromone signalling itself. Remarkably, none of the tested strains abrogated habituation altogether. However, their habituation time varied a lot (Figure 5B, Supplementary Figure 6). Whereas deletion of the control genes failed to affect habituation behaviour compared to the wild type, thirteen of the hit strains habituated faster and thirteen slower than wild type control. Only 6 hit strains showed no significant change in habituation time. The most extreme phenotype was observed in the *nat1Δ* mutant cells, which habituated extremely fast (median response time of 45 min). The Nat1 protein is part of a conserved N-acetyl transferase complex and functions in modification of factors involved in DNA silencing and cell cycle regulation, among others. Importantly, deletion of courtship genes typically caused rapid habituation, whereas deleting habituation genes delayed habituation. Thus, in most cases the phenotypic and screening data were fully consistent with each other. The only exceptions were *cln3*Δ, *cik1*Δ and *ydr102c*Δ cells, which changed habituation time in the opposite direction as predicted. Indeed, *CLN3* scored as a courtship gene because insertions in the PEST sequence are strongly selected in pheromone. However, removal of the whole gene abrogates the advantageous effect of these insertions and strongly delays habituation. *CIK1* codes for a microtubule motor subunit of the kinesin complex and functions in nuclear migration and karyogamy during mating. While it was found to be a sporadic courtship gene in the screen, deletion mildly but significantly delayed habituation. *YDR102C* is an uncharacterized ORF scoring as courtship gene in the SATAY screen, i.e., inhibiting habituation. However, its full deletion strongly delayed habituation. As it is located upstream of *STE5*, a key pheromone signalling gene, its deletion or transposon insertion within the gene might affect *STE5* expression in opposite ways.

In the wild type, the time cells needed to habituate increased with increasing pheromone concentration (Figure 1B). Few mutants failed to modulate their habituation behaviour as a function of pheromone concentration. This included the *kar4Δ* mutant cells (Figure 5C, Supplementary Figure 5B), which showed the same response time over many concentrations, unlike wild type cells. Thus, the transcription and RNA-editing factor Kar4 helps encoding information about pheromone concentration. At least one more gene, *RPN4*, showed a similar phenotype (Supplementary Figure 5C). Rpn4 is a transcription factor inducing proteasome synthesis in response to stress and was one of the mutants that initially failed to show a change in habituation time.

Independently of pheromone concentration, wild type cells maintained their habituated state and ignored the pheromone for many hours and division cycles after resuming proliferation (Figure 5D). Whereas only 1-8% of the wild type cells re-shmooed at least once within the imaging time (18 hours), 6 (nearly 20%) of the tested hits showed a substantial increase in re-shmooing frequency (Figure 5E). This included the *mum2Δ* mutant cells, which otherwise showed no other habituation phenotypes under our experimental conditions. Interestingly, for each of these six mutant strains, only up to 50% of cells in the population re-responded to pheromone at least once. Thus, memory is distributed over the network and not stored in a single factor.

Another striking phenotype concerned the pheromone response of daughter cells born by habituated mother cells. In the wild type, these daughters are born naïve and respond to pheromone normally. Therefore, the median response time of wild type daughter cells was only slightly reduced compared to that of their respective mother cells (Figure 5F, Supplementary Figure 5D). In contrast, the daughter cells of *far11Δ* and *rpn4Δ* mutant cells habituated much faster than their mothers. Especially the *rpn4Δ* mutant daughter cells barely responded to the pheromone signal, indicating that they inherited memory from their mother cell.

Of the 32 mutant strains tested, only four did not show a noticeable change in any of the monitored phenotypes compared to the wild type (*UBR2*, *DIG1*, *YAP6*, *CPR7*), although two of them (*UBR2* and *DIG1*) were consistent hits. The *YAP6* and *CPR7* genes were selected for their sporadic behaviour, with *CPR7* passing cutoff in two out of the three original screens. These 4 genes might influence habituation under conditions different than those we tested here. Their lack of effects in our experiments may also reflect the sporadic nature of many hits. Remarkably, 12 out of the 14 genes selected for being replicate-specific hits showed at least one clear habituation phenotype. Thus, the genes showing sporadic phenotypes are indeed bona fide members of the habituation network. The fact that they were variable in the screen, i.e., at low pheromone levels, and reproducible in the phenotypic analysis may indicate that sporadicity is stronger at physiological pheromone levels. Furthermore, these phenotypic analyses demonstrated that the components of this network control a broad range of response parameters, sometimes several of them. Thus, habituation entails collecting, deconvolving, encoding and utilizing a broad range of information about the cell’s history and environment to determine whether responding to a pheromone signal is meaningful or not. Furthermore, the fact that habituation was never abrogated all together in any of the tested mutants suggests that it is an emergent property of a collective of genes and their products rather than an outcome ultimately controlled by few dedicated genes. In other words, the robustness of the habituation process highlights the redundancy of the habituation network.

## Discussion

Here, we extensively describe for the first time the genetic network underlying a simple, habituation-like, learning task. This network is surprisingly large (at least 5% of the genome) and broadly distributed. Under physiological conditions, many network components acted only sporadically (at least 68% of hits), indicating that the weights of most nodes vary. Thus, this distributed network is both highly redundant and intrinsically plastic. Both, our screens in mutant strains and the phenotypic analysis performed with strains lacking individual network genes demonstrate that habituation is a highly robust process. Together, these data establish that habituation is a plastic process supported by the emergent properties of a large gene network, rather than a hard-wired, pre-scripted procedure. The fact that this network supports habituation at the single cell level allows making the following points about cellular learning.

First, the implication of many cellular functions in the habituation network suggests that it integrates a rich body of information, including external and internal signals. This suggests that habituation to a single stimulus such as pheromone engages more than that stimulus alone. Supporting this idea, previous work has established that the response of yeast cells to pheromone is strongly influenced by the condition of the individual cells, such as their age^26–28^, and external context, such as nutrient availability^29^ and the presence of stressors^30^. At least in the case age, reduced response is due to habituation being accelerated in old cells^26^. Thus, we speculate that this will be similar in many other contexts and that some of the functionalities represented in the habituation network funnel this contextual information into the habituation process. In turn, the richness of the network also suggests that it may support habituation to other stimuli as well. Likewise, this network might support other learning processes, such as sensitization or associative learning. Overall, our observations suggest that single cells might have a much richer spectrum of behavioural capacities than generally presumed, and that the learning capacities of yeast cells naturally emerge from the interconnection of cellular pathways in a large genetic network. As such, learning capacities are likely to emerge similarly in any other cell type, where cellular pathways are equally likely to be interconnected.

Second, and logically deriving from the first point, our data suggest that cellular learning is supported by potentially sophisticated computational capacities born by the identified gene network. The fact that inactivating distinct single network components affected different aspects of habituation, like timing, concentration-dependency and memory, suggests that this network comprises subnetworks processing specific elements of information. For example, a substructure comprising the transcription and mRNA editing factor Kar4 and the transcription factor Rpn4 extracts and propagates information about pheromone levels. Also, genes acting in very different parts of the network contributed to memory, suggesting that they might store different information components. Thus, pheromone habituation offers a rich paradigm for studying how cells decompose information, store it and use it to make appropriate decisions.

Third, the fact that many nodes act only sporadically rather than consistently indicates that their weight is not constant. Although some of the lack of consistency highlighted by the replicate-to-replicate variability might reflect a lack of statistical power, as demonstrated when including the results of the epistasis screens, a large fraction of this variability appears to be intrinsic. Indeed, the power of our approach is not only to identify hits, but to also unambiguously pin-point neutral genes. This demonstrated that many sporadic hits are truly behaving neutrally in some of the replicates. On one hand, this variability might reflect differences in contextual conditions between the replicates, such as growth and processing conditions. This would give different weight to the genes distinctly influenced by these contextual differences. Indeed, the three wild type replicates were carried out at different times. However, to the best of our capacity, the immediate screening conditions were kept extremely consistent. Thus, this replicate-to-replicate variations may indicate that the habituation network is extremely sensitive to small environmental changes. If so, it will be interesting to investigate what the functional relevance of such a sensitivity is. However, replicate-to-replicate variability might on the other hand reflect differences in the state of the habituation network in distinct populations. If the habituation network enables cells to memorize their experience and learn, as suggested by our results, a prediction is that at least some of the nodes change weight as a function of the cells’ history. As a consequence, their signature in our screen may change. However, to be detectable, these changes must have an impact at the population level. This could be the case if the cells communicate with each other, for example through metabolites, or if network states can be stably propagated across generations. The latter could be achieved through prion conversion^19,31^ or epigenetic changes at gene levels (affecting single nodes) or self-amplifying feedback loops (affecting network motifs)^32,33^. Supporting this idea, the network is rich in such motifs, including epigenetic players such as the Set1/COMPASS methyltransferase and Mediator complex^34–36^, and proteins with prion-like domains (at least 26, 9% of the hits). Proteins with prion-like domains may propagate information in their conformational state^37^. For example, we previously showed that the prion-like Q- and N-rich domains of the RNA-binding protein Whi3 contribute to memory of pheromone exposure^19,26^. Importantly, however, three of the six genes identified to affect memory in our phenotypic studies have neither Q- or N-rich domains nor are involved in chromatin. These observations suggest that other memory mechanisms may still be discovered.

In summary, our study establishes that at physiological conditions, the cells’ response to input signals is highly plastic, as is the underlying genetic network. Therefore, we may have to move away from the notion that cells respond in algorithmic, pre-determined manners implemented by hard-wired, hierarchical, linear pathways. We speculate that classical response pathways represent the state of a regulatory network when one signal strongly dominates. In the future, the challenge will be to develop new approaches and concepts for studying large plastic networks, how they respond to dynamic environments and how they bring temporal depth to cellular behaviour. For example, we anticipate that our studies still underestimate the size of the habituation network that we describe here. Repeating our screen under a broad range of environmental conditions is likely to identify other subnetworks that become important in specific contexts.

In recent years, reproducibility of results has become an important issue that keeps on sparking discussions in the scientific community and beyond. Here, we show that even when experiments are repeated in highly controlled manners, the results show an intrinsic lack of reproducibility. The data suggest that non-reproducibility is not necessarily a sign of noise or that experiments were not properly carried out. It can rather be indicative that the process of interest is investigated near physiological conditions and modulated by a large regulatory network with deep adaptive capabilities. It should encourage us to reconsider data that seem irreproducible, as they might not be variable by random but depict a more complex system able to incorporate contextual information and past experiences into decision making, i.e., to show learning behaviours.

## Material and Methods

### Yeast strain generation

All yeast strains used in this study are based on *S. cerevisiae* BY4741^38^ and are listed in Supplementary File 1. Strains were generated by crossing or transformation of marker genes flanked by homologous sequences to the target locus using an optimized lithium acetate/ single-stranded DNA/ PEG transformation protocol^39^. After transformation, clones were selected on yeast extract peptone dextrose (YPD) agar plates supplemented with nourseothricin (100 µg/ml), geneticin (500 µg/ml) or hygromycin (300 µg/ml) or on synthetic complete (SC) drop-out plates and correct gene deletion was verified by PCR. Yeast strains were cultured in standard YPD or SC medium with 2 % glucose with shaking at 160 rpm and stored at-80 °C in YPD with 30 % glycerol.

### SATAY screen

For transposon library generation, strains were transformed with the pBK549 plasmid provided by the Kornmann lab, which encodes the MiniDs transposon within the ADE2 gene, a hyperactive Ac transposase under control of the *GAL1* promoter and *URA3* as selective marker^21^. Single transformed colonies were grown in pre-induction medium (SD complete-uracil +2% raffinose +0.2% glucose) over night before cells were spread on at least fifty induction plates (SD complete -adenine +2% galactose) per library to induce transposition. After three weeks of incubation at 30 °C, libraries were harvested, diluted to OD600 of 0.08 in 1 liter of SD -adenine +2% galactose medium and grown to saturation overnight. For selection, libraries were inoculated into 500 ml YPD +20 µg/ml casein and 0, 4, 6 or 8 nM α-factor (GenScript) at OD600 of 0.1 and grown for 24 hours. This step was repeated before cells were harvested for genomic DNA extraction and sequencing.

Genomic DNA was extracted using phenol-chloroform-isoamyl alcohol following the published SATAY protocol^21^. For sequencing, each library was digested with DpnII and NlaIII (NEB) separately before circularization of the DNA using T4 DNA ligase (Thermo Scientific) and amplification of the transposon-insertion sites by PCR (Taq polymerase, NEB) using a universal primer (P5_MiniDs) combined with individual MiniDs_P7 index primers for multiplexing of 24 samples (Supplementary File 2). Libraries were sequenced on an Illumina NextSeq 500 instrument using the NextSeq 500/550 High Output Kit v2.5 (75 Cycles) following the manufacturer’s protocol (Illumina).

### SATAY data analysis

The resulting raw reads were cleaned by removing adaptor sequences, low-quality-end trimming, and removal of low-quality reads using BBTools v. 38.18 [Bushnell, B. BBMap. Available from: https://sourceforge.net/projects/bbmap/]. The exact commands used for quality control can be found on the Methods in Microbiomics webpage (Sunagawa, S. Data Preprocessing-Methods in Microbiomics 0.0.1 documentation. Available from: https://doi.org/10.5281/zenodo.15019381).

Clean SATAY alignment files were processed using a custom satay_tools package [https://github.com/MicrobiologyETHZ/satay_tools] to identify transposon insertion sites and quantify insertion frequencies across genomic features. The processing workflow consisted of the following steps:

Read alignment: The quality-controlled reads are aligned against *Saccharomyces cerevisiae* S288C reference genome R64 (GCF_000146045.2) using STAR v. 2.7.8 aligner^40^.

BAM File Processing: Multiple BAM files from DpnII and NlaIII libraries are merged using samtools v. 1.22.1 ‘samtools merge’ to create consolidated alignment files for each biological sample^41^. Quality filtering is then applied using ‘samtools view’ with parameters-F 256,272 -q 10 to exclude secondary alignments (reads failing quality checks and reads with mapping quality scores below 10. Filtered alignments are converted to BED format using bedtools v. 2.31.1 ‘bamtobed’ for downstream processing^42^.

Insertion Site Determination: Precise transposon insertion coordinates were determined by analysing read orientation. For reads mapping to the positive strand, the insertion site was defined as the read start position (5’ end), while for reads on the negative strand, the insertion site was defined as the read end position (3’ end).

Data Consolidation and Filtering: Insertion sites were sorted by genomic coordinates using standard Unix sort utilities, then merged using ‘bedtools merg’ with strand-specific consolidation (-s flag) to combine multiple reads mapping to identical insertion sites. The merge operation counted the number of supporting reads per insertion site and preserved strand information. Low-confidence insertion events supported by only a single read were filtered out to reduce noise and focus analysis on high-confidence integration sites.

Genomic Feature Mapping: Filtered insertion sites were mapped to genomic features of interest using ‘bedtools map’. For each genomic interval (e.g. gene), two metrics were calculated: (1) the total number of unique insertion sites within the interval, and (2) the total number of reads mapping to the interval. This dual quantification provides both insertion density and read depth information for downstream fitness analysis.

Cross-Sample Integration: Results from individual samples were consolidated into count matrices using pandas data processing. For GFF-formatted annotations, gene identifiers and genomic coordinates were extracted. The final output consisted of two matrices: transposon insertion counts (number of unique insertions per gene per sample) and read counts (total supporting reads per gene per sample).

Statistical analysis: Statistical testing of fitness effects was performed using PyDESeq2^43^, a Python re-implementation of the DESeq2 framework^44^. Read- or insertion-count matrices were filtered to retain only genes with a total count above 20 unique insertions / 200 reads across all samples. The statistically significant hits were then identified using Wald test with Benjamini–Hochberg multiple testing correction.

Single replicate analysis: For each replicate, raw counts (either insertion or reads counts) were library-size normalized and log-transformed using a TPM-like transformation, log2(cij / Σi cij × 106 + 0.5), where cij is the count for gene i in sample j and 0.5 is a pseudocount to stabilize zeros. To rank the genes, per-gene log2 fold changes were then computed for each replicate-matched pair (treated vs. 0 nM α-factor of the same replicate) as the difference of normalized values. Genes with insufficient insertions/reads were excluded from the ranking. Specifically, genes with a paired raw-count sum below 20 (transposon analysis) or 200 (read analysis) were masked, and Z-scores were computed only across the genes passing the filter, separately for each replicate-pair fold-change distribution.

For each gene, the more extreme z-score (single replicate analysis) or lower adjusted p-value (statistical analysis) obtained from comparing the number of insertions per gene and reads per gene was used for all analyses shown throughout the paper.

### Yeast cultivation and microscopy

For analysis of pheromone response and habituation, yeast strains were grown to exponential phase in SD complete overnight, diluted to OD600 of 0.05 in fresh SD complete in the morning and again grown to OD600 between 0.2 and 0.4. For imaging, cultures were diluted to OD600 0.18 in SD complete before 5 µl of cells were pipetted into the wells of an 8-well microscopy slide (µ-slide 8-well glass bottom slides, Ibidi) and were covered with a piece of fresh SD complete agar containing α-factor (GenScript) not older than 6 days). For the analysis of memory inheritance, cells were diluted to OD600 0.05 and loaded onto a commercially available CellASIC® ONIX Y04C-02 microfluidic plate for haploid yeast cells (Merck Millipore). For imaging, cells were continuously flushed with SD complete medium containing α-factor using the CellASIC® ONIX2 Microfluidic System at a pressure of 2 psi.

Timelapse microscopy was performed on an inverted Nikon Eclipse TiE microscope (Nikon Instruments) using a hardware-based automated focusing system (Perfect Focus System (PFS)), a motorized XY-stage, a CoolLED pE-300 light source for fluorescence imaging (CoolLED), and a Plan Apo 60 × 1.4 NA oil immersion objective (Nikon Instruments). The temperature of the incubation chamber was set to 25 °C. We acquired 15-20 brightfield and fluorescence images for each strain at a single focal plane every 15 min for 18 hours. Imaging parameters, filters and fields-of-view to image were controlled using the µManager software^45^.

### Microscopy data analysis

Image analysis was done with ImageJ Fiji^46^. Cell cycle arrest, time to habituation and repeated responses were scored manually based on localization of the cell cycle marker Whi5-tdTomato and cell morphology and recorded using a macro in ImageJ Fiji (Available from: https://github.com/faultby/my-fiji-scripts/tree/main). A cell was counted as arrested in G1 when the red signal was clearly visible in the nucleus, while the first frame without a nuclear Whi5-tdTomato signal was counted as time of habituation. A prolonged nuclear Whi5-tdTomato signal after initial return to budding for at least 60 min was counted as re-shmooing. For each strain and condition, at least fifty individual cells were analysed. Median times to habituation were normalized to the wild type strain imaged in the same experiment. Differences in the distribution of response times between the strains and the wild type were compared using a two-sided Mann–Whitney U test with Benjamini–Hochberg correction applied across all comparisons.

### Protein-protein interaction networks

Screening results were filtered to use all genes with an adjusted p-value ≤0.05 (combined statistical analysis) or z-score ≤-3 or ≥+3 (individual replicates) as input for multiple protein interaction analysis using STRING^23^. Predicted interactions above an interaction score of 0.4 (medium confidence) were exported. Networks were visualized and manually clustered by function using Cytoscape^47^.

### Resource availability

Requests for further information and resources should be directed to and will be fulfilled by the lead contact, Yves Barral. All yeast strains generated in this study are available upon request. The original imaging data and data analysis files have been deposited in the ETH Research Collection (https://doi.org/10.3929/ethz-c-000800474). Genomic data are available in the European Nucleotide Archive under the accession number PRJEB113239. The code used in this publication is described in the Methods and is available from https://github.com/MicrobiologyETHZ/satay_tools.

### Author contributions

L.H. designed and performed experiments, analysed the results of screens and validation experiments and visualized the data. I.P. preformed the SATAY screens. A.D. contributed to yeast strain generation and phenotypic characterization of the mutants. A.M. and B.K. developed the SATAY protocol and provided support during the screens. A.S. set up the data analysis pipeline and performed the sequencing data analysis. L.H. and Y.B. conceptualized the project and drafted the manuscript. Y.B. acquired the funding.

## Supplementary figures

**Supplementary figure 1:**
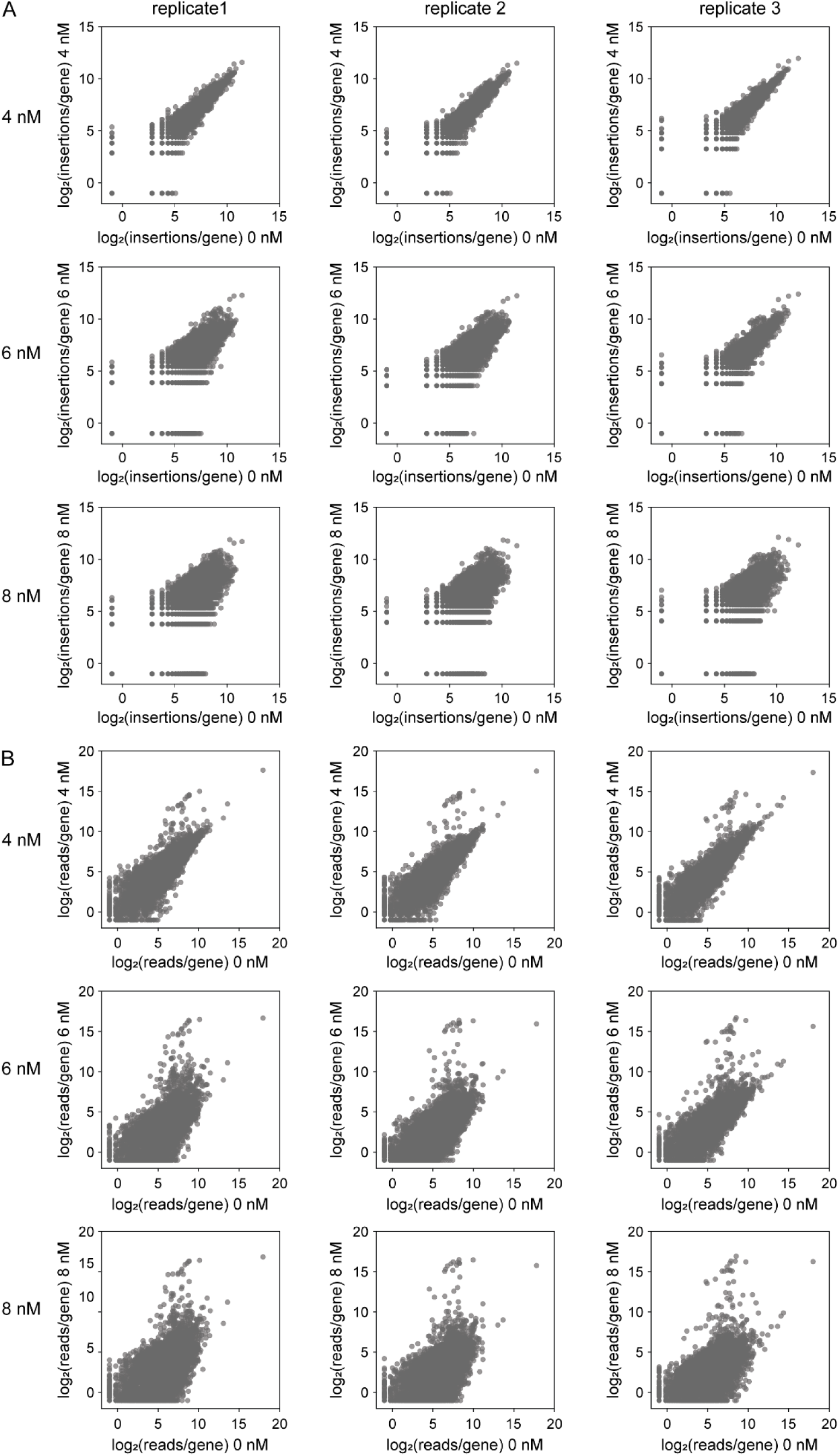
Normalized numbers of transposon insertions and reads per gene for all three replicates for the libraries selected at 4, 6 and 8 nM pheromone compared to the same library at 0 nM.

**Supplementary figure 2:**
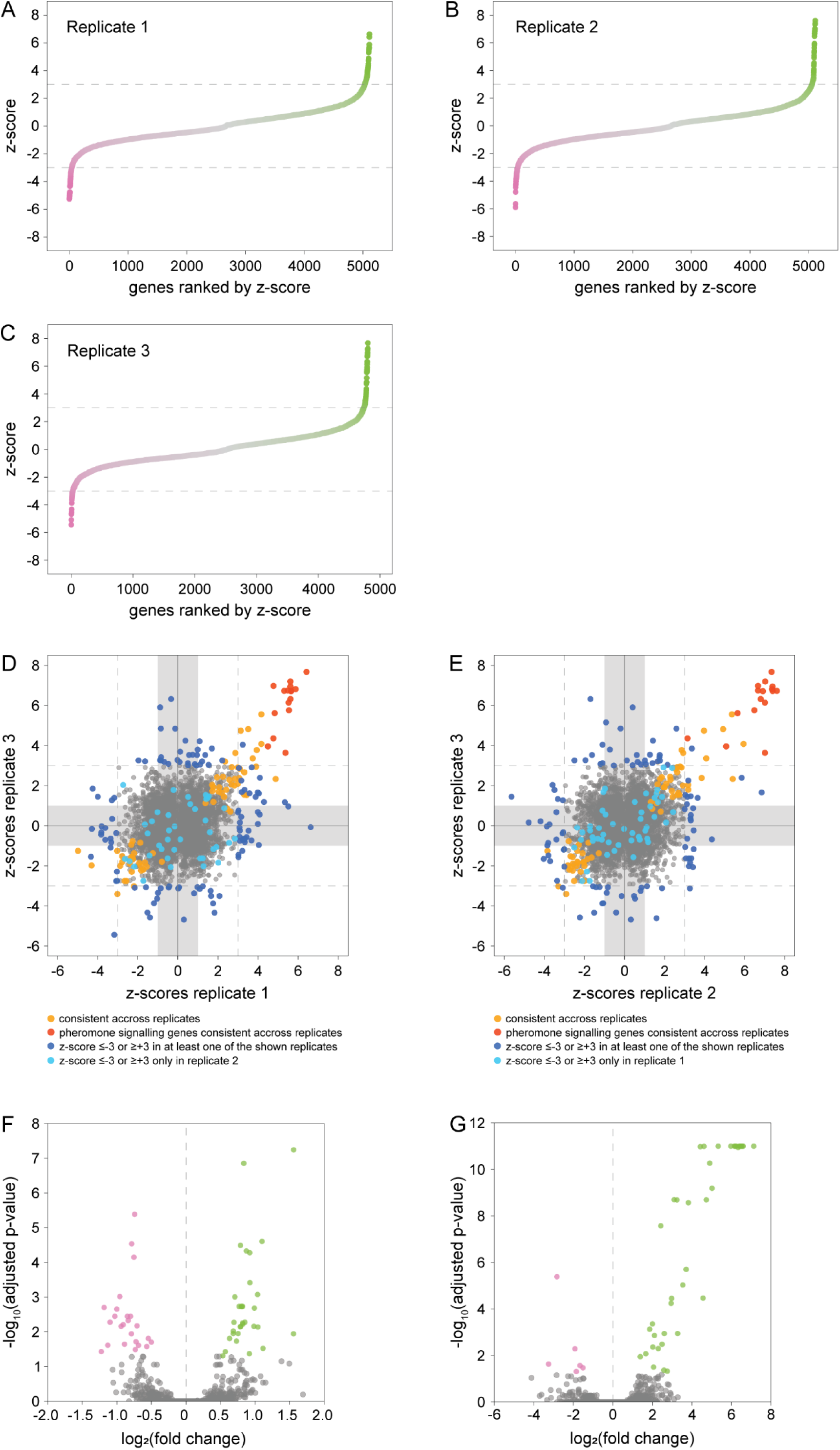
**A-C)** Genes ranked by their z-score for the three individual replicates of the SATAY screen performed at 0 vs. 4 nM pheromone. **D)+E)** All genes with a measurable z-score in replicates 1 and 3 or replicates 2 and 3 are plotted as a function of their z-scores. Grey area: z-score values within one standard deviation. Dark blue dots: genes that pass the z-score threshold of ≤-3 or ≥+3 in at least one of the two replicates. Light blue dots: genes that pass the threshold of ≤-3 or ≥+3 only in the third replicate. Orange dots: statistically significant (consistent) genes across all three replicates. Red dots: Expected pheromone signalling genes consistent across all three replicates. Grey dots: Neutral in each replicate and across the three replicates. **F)** Fold changes and adjusted p-values of all genes identified in the statistical analysis across all three replicate screens calculated based on the number of transposon insertions per gene at 0 and 4 nM pheromone. Magenta dots: consistent habituation gene (depleted of insertions), Green dots: consistent courtship or pheromone signalling gene (enriched in insertions), cutoff: adjusted p-value ≤0.05. **G)** Fold changes and adjusted p-values of all genes identified in the statistical analysis across all three replicate screens calculated based on the number of sequencing reads per gene at 0 and 4 nM pheromone. Colours and cutoff as in F).

**Supplementary figure 3:**
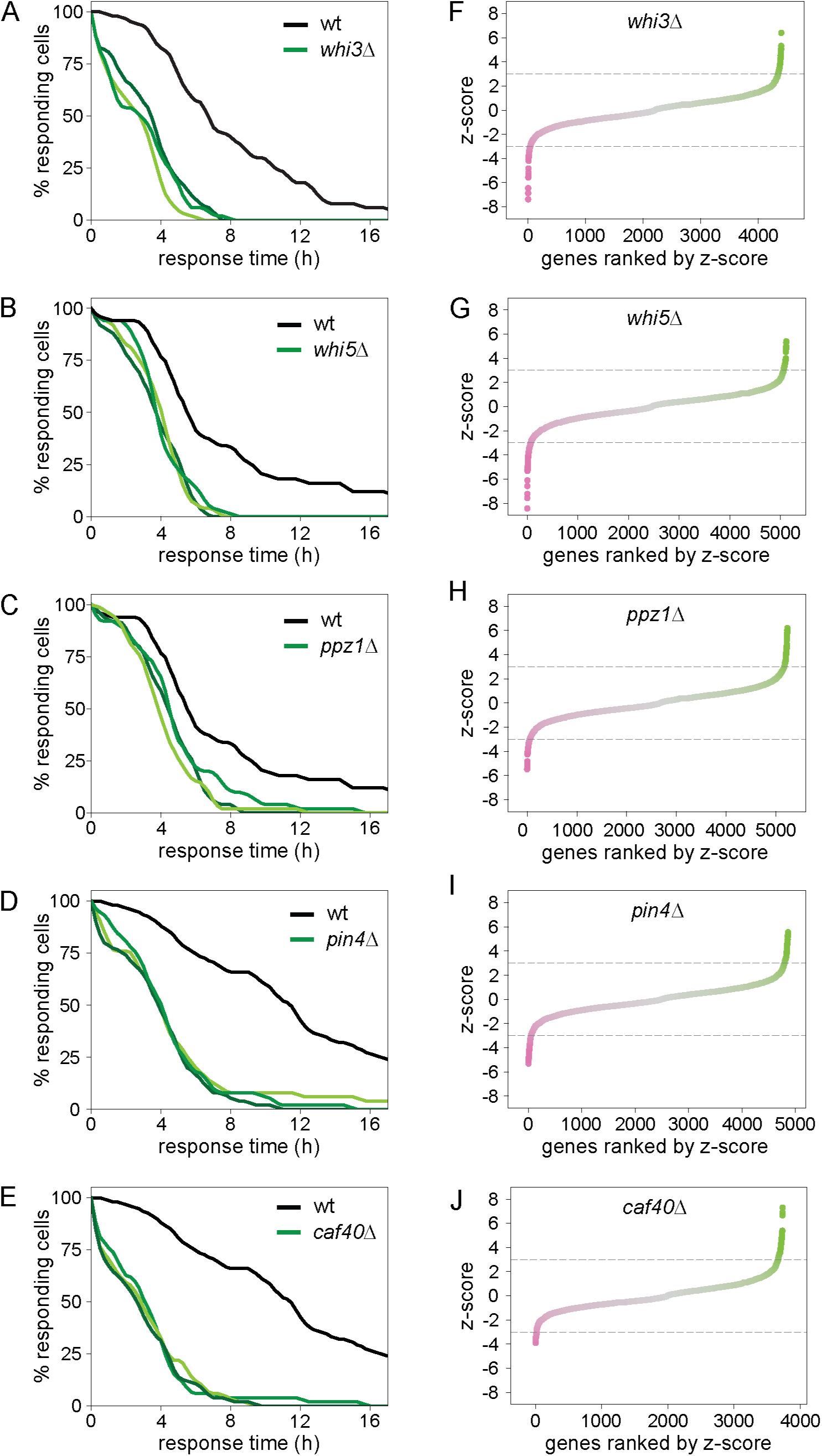
**A)-E)** Time to habituation of individual query gene deletion strains in response to 12 nM α-factor as measured by time-laps microscopy. For each mutant, three biological clones were analysed. The wild type control imaged in the same experiment is shown in all panels. Data for 50 cells per sample. **F)-J)** Genes ranked by their z-score from the SATAY screens of the individual query gene deletion strains performed at 0 vs. 8 nM pheromone.

**Supplementary figure 4:**
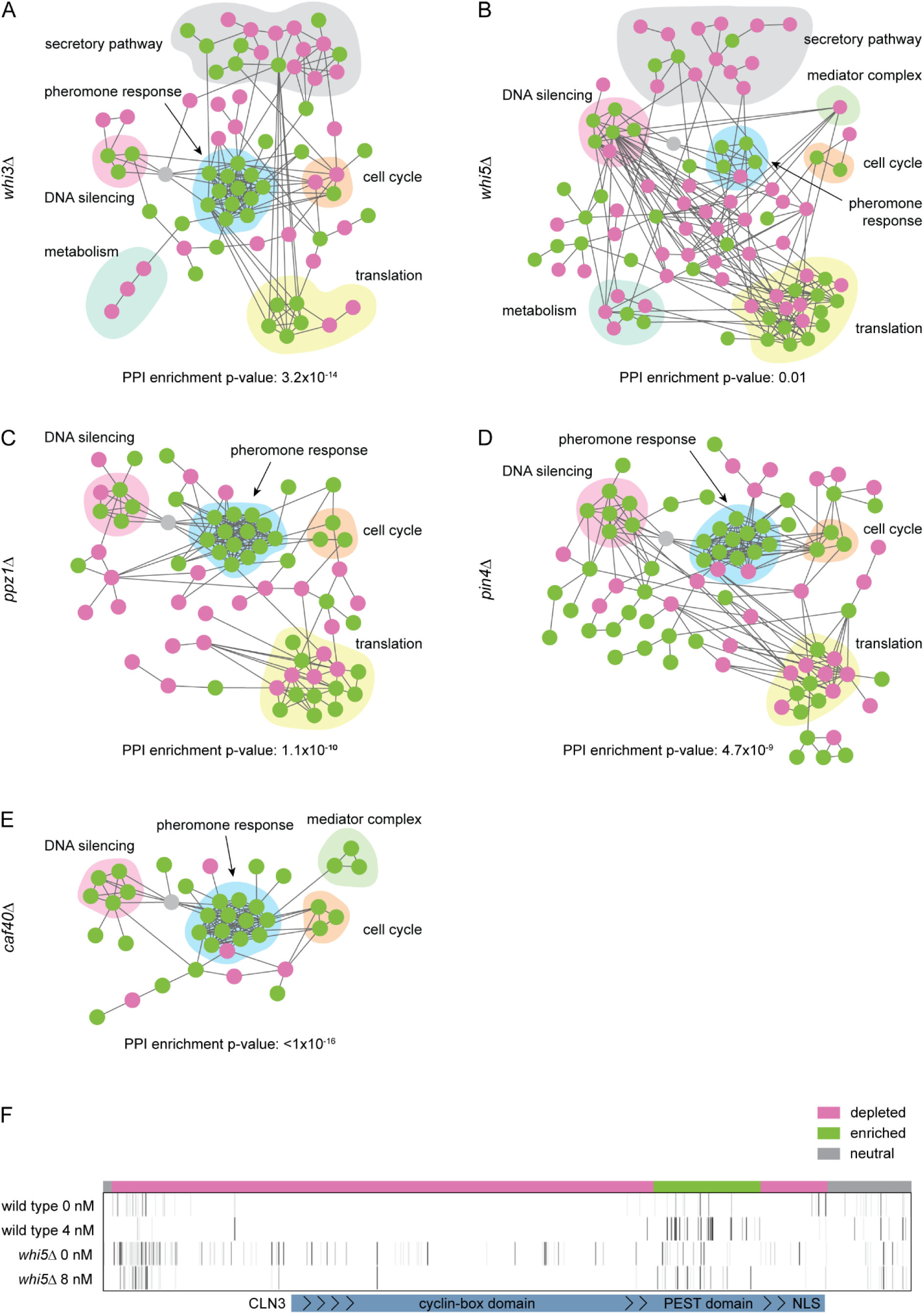
**A)-E)** Predicted protein-protein interaction network for the genes with z-score ≤-3 or ≥+3 in the individual query gene deletion strains (*whi3*Δ: 94, *whi5*Δ: 126, *ppz1*Δ: 116, *pin4*Δ: 115, *caf40*Δ: 86). *HML*α (grey) is used as an anchor. Only the main fully connected networks are shown (*whi3*Δ: 70, *whi5*Δ: 92, *ppz1*Δ: 68, *pin4*Δ: 84, *caf40*Δ: 39). Courtship and pheromone signalling genes are plotted in green, habituation genes in magenta. **F)** Mapped transposon insertions (vertical lines) in the *CLN3* genomic region in the wild type and *whi5*Δ strains. Comparison of the insertion pattern in the presence and absence of pheromone identifies domain specific differences in number and distribution of insertions between the strains and conditions.

**Supplementary figure 5:**
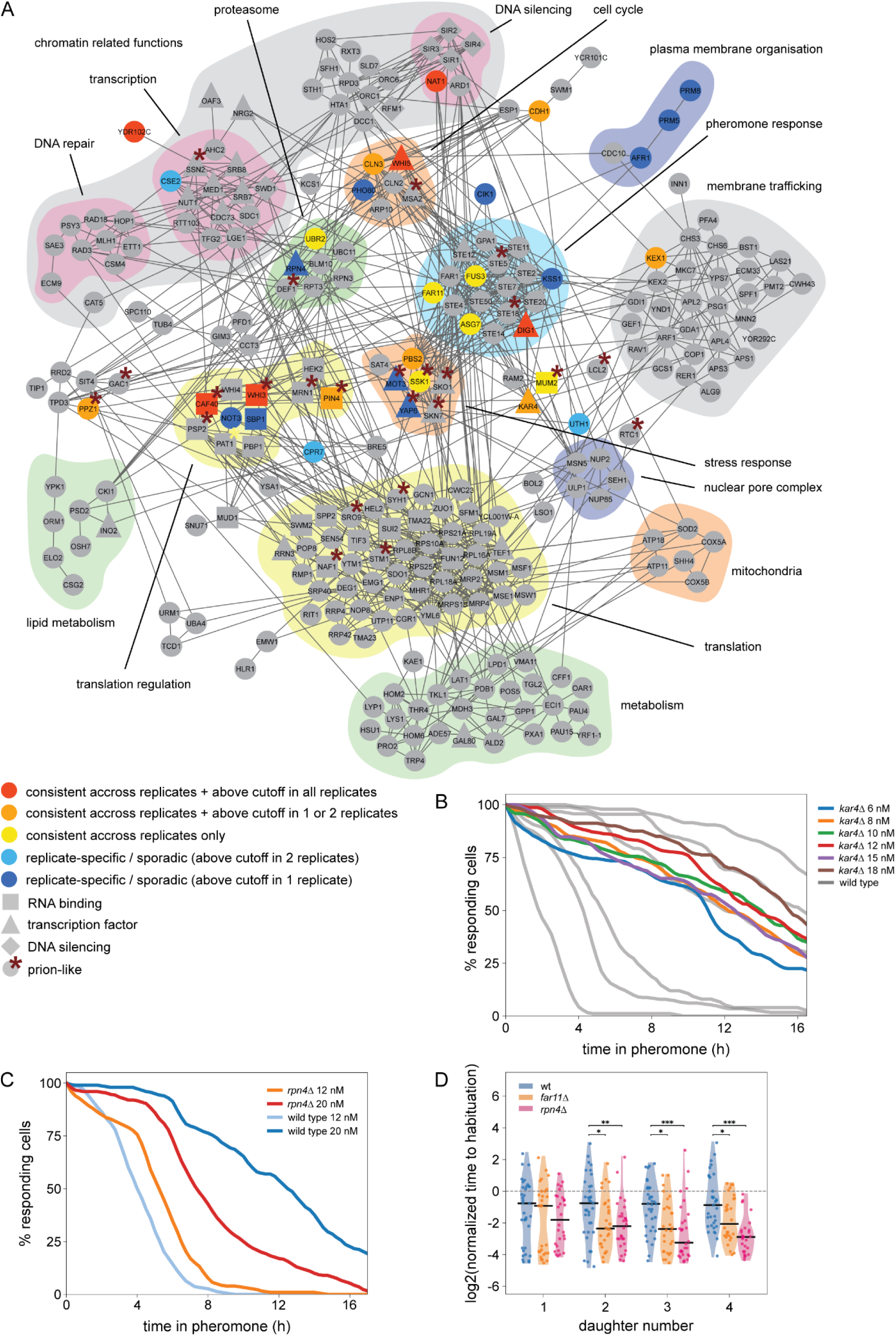
**A)** Predicted protein-protein interaction network of all hits identified in the individual screens (at cutoff of z≤-3 or ≥+3) and the combined statistical analysis (adjusted p-value ≤0.05). 294 hits in total resulted in a network of 251 connected nodes using STRING (enrichment p-value <10^-16^). Five additional genes which were just below read / transposon number abundance cutoff (*PRM5*, *SBP1*, *NOT3, CSE2*) or only identified as a hit in the *whi5*Δ strain (*KSS1*) and affected habituation in our validation experiments (see Figure 5B) were also included. Hits were manually ascribed to functional clusters. Node colour and shape code is indicated within the figure. All coloured nodes were selected for gene deletion and phenotypic analysis. **B)** Time to habituation in *kar4Δ* mutant cells at indicated α-factor concentrations. The duration of shmooing was determined by time-laps microscopy. Grey curves represent wild type under the same conditions (see Figure 1B) (wild type: 6 nM: 85 cells, 8 nM: 114 cells, 10 nM: 50 cells, 12 nM: 82 cells, 15 nM: 88 cells, 18 nM: 94 cells; kar4Δ: 6 nM: 98 cells, 8 nM: 80 cells, 10 nM: 98 cells, 12 nM: 150 cells, 15 nM 66 cells, 18 nM: 80 cells). **C)** Time to habituation in wild type and *rpn4Δ* mutant cells at indicated α-factor concentrations. The duration of shmooing was determined by time-laps microscopy. Data from 50 individual cells per sample are shown. **D)** Habituation time in daughter cells born by habituated wild type, *far11Δ* and *rpn4Δ* mutant mother cells. Shmooing duration of the mother cells and their first four daughter cells was determined by time-laps microscopy in the presence of 12 nM α-factor. Habituation times of daughter cells were normalized to their respective mother cell. Data points represent individual daughter cells (27 to 35 cells per strain), black lines represent median values. Differences in normalized time to habituation between strains were analysed using a two-sided Mann–Whitney U test (* p-value <0.05, ** p-value <0.01, *** p-value <0.001).

**Supplementary figure 6:**
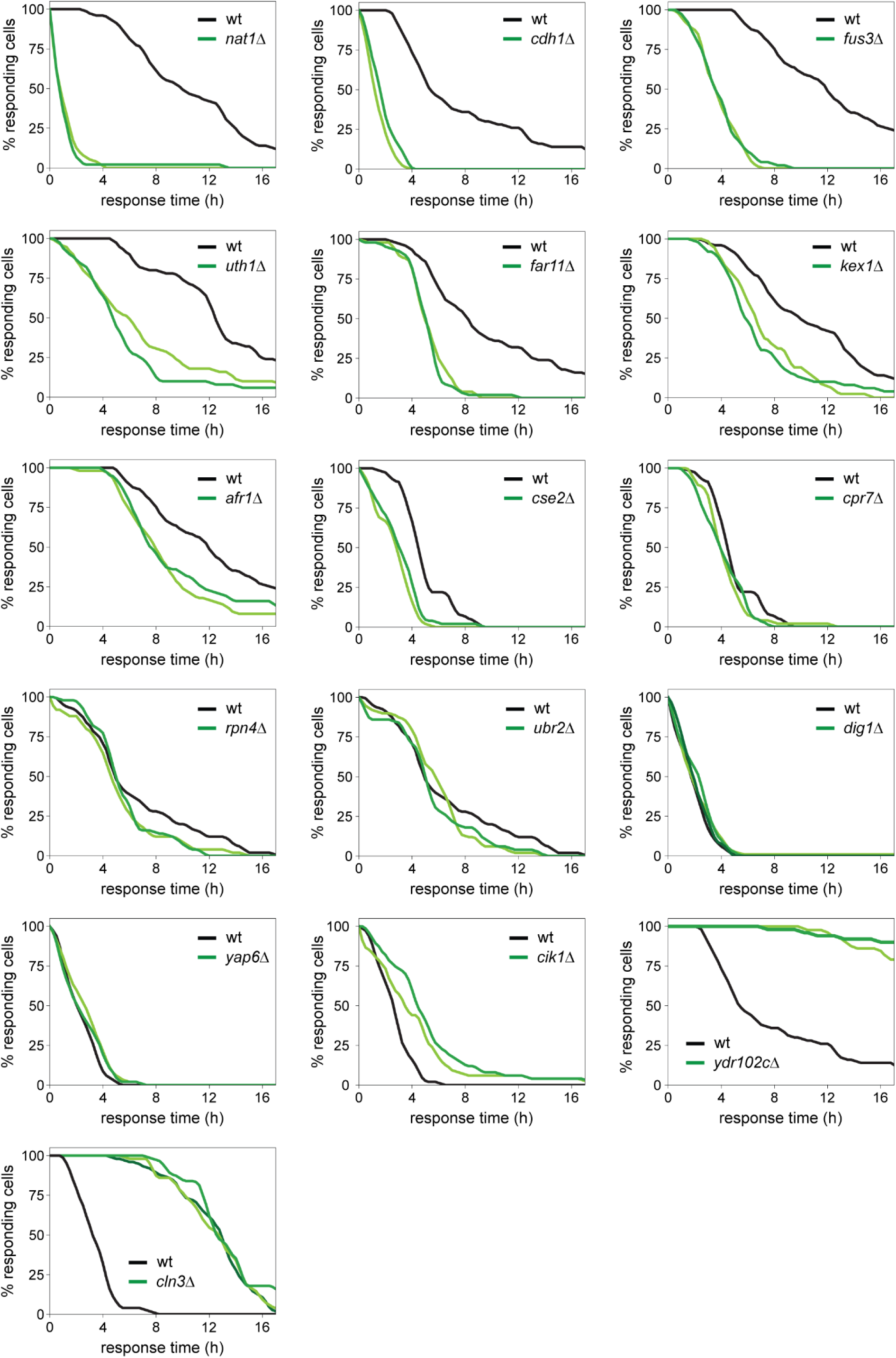
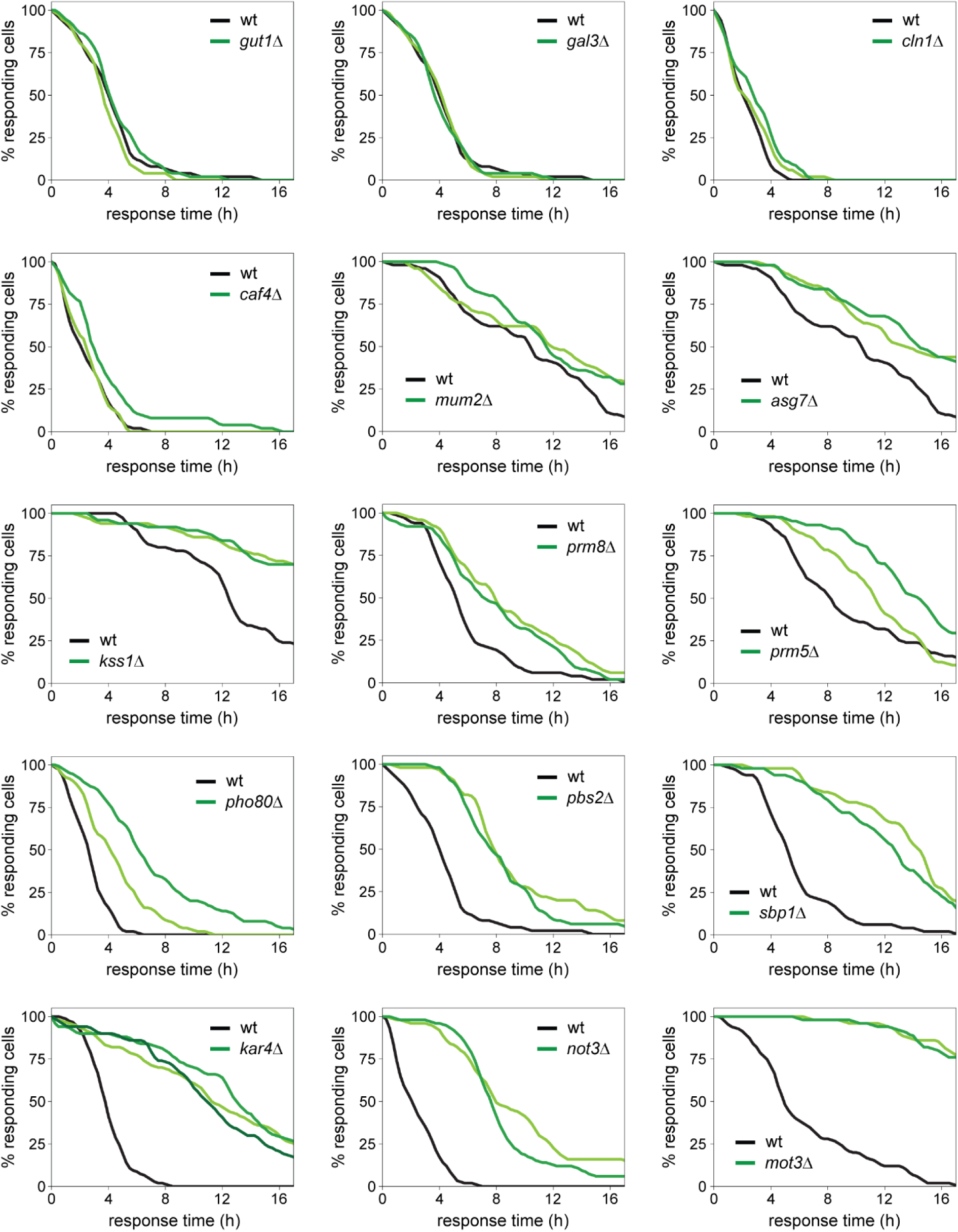
Time to habituation of individual gene deletion strains in response to 12 nM α-factor as measured by time-laps microscopy. For each mutant, at least two biological clones were analysed. The wild type control imaged in the same experiment is shown in all panels. Data for 50 cells per sample.

